# Emx1-Cre is expressed in peripheral autonomic ganglia that regulate central cardiorespiratory functions

**DOI:** 10.1101/2022.03.03.482724

**Authors:** Yao Ning, Jeffrey L. Noebels, Isamu Aiba

## Abstract

The Emx1-IRES-Cre transgenic mouse is commonly used to direct genetic recombination in forebrain excitatory neurons. However, the original study reported that Emx1-Cre is also expressed embryonically in peripheral autonomic ganglia, which could potentially affect the interpretation of targeted circuitry contributing to systemic phenotypes. Here, we report that Emx1-Cre is expressed in the afferent vagus nerve system involved in autonomic cardiorespiratory regulatory pathways. Our imaging studies revealed expression of Emx1-Cre driven tdtomato fluorescence in the afferent vagus nerve innervating the dorsal medulla of brainstem, cell bodies in the nodose ganglion, and their potential target structures at the carotid bifurcation such as the carotid sinus and the superior cervical ganglion. Photostimulation of the afferent terminals in the nucleus tractus solitarius (NTS) *in vitro* using Emx1-Cre driven ChR2 reliably evoked excitatory postsynaptic currents in the postsynaptic neurons with electrophysiological characteristics consistent with the vagus afferent nerves. In addition, optogenetic stimulation targeting the Emx1-Cre expressing structures identified in this study, such as vagus nerve, carotid bifurcation, and the dorsal medulla surface transiently depressed cardiorespiratory rate in urethane anesthetized mice *in vivo*. Together, our study demonstrates that Emx1-IRES-Cre is expressed in the key peripheral autonomic nerve system and can modulate the cardiorespiratory function independently of forebrain expression. These results raise caution when interpreting the systemic phenotypes of Emx1-IRES-Cre conditional recombinant mice, and also suggest the utility of this line to investigate the modulators of afferent vagal system.

**Significance Statement:** Emx1-IRES-Cre mice are widely used to dissect critical circuitry underlying neurological disorders such as epilepsy. These studies often assume the Cre is expressed selectively in forebrain excitatory neurons. However, earlier work reported that Emx1 is expressed in several peripheral tissues of the developing embryo and thus gene recombination may affect these peripheral structures. In this study, we characterized the expression and physiological functions of Emx1-Cre expressed in the vagus nerve, the critical peripheral autonomic component. Optogenetic stimulation of these Emx1-Cre^+^ neurons activates the nucleus tractus solitarius neurons within the brainstem *in vitro* and induces complex cardiorespiratory reflex *in vivo*. This study confirmed that peripheral Emx1-Cre^+^ cells are involved in autonomic regulation and potentially affect transgenic mouse phenotypes.

## Introduction

Emx1 is a homeobox gene widely expressed in excitatory neurons in the mammalian forebrain during embryogenesis, including the cerebral cortex, hippocampus, and olfactory bulbs (Simeone et al., 1992; Gorski et al., 2002). A mouse line carrying Cre recombinase under the endogenous Emx1 promoter (i.e. Emx1-IRES-Cre (Gorski et al., 2002)) has been widely used in studies of neurological diseases to elucidate the functions of specific genes in these neurons. In epilepsy studies, developmental Cre-loxP recombination strategy (Orban et al., 1992) has been contributing to the identification of many pathogenic mechanisms of genetic epilepsy (Boillot et al., 2014; Soh et al., 2014; Makinson et al., 2017; Klofas et al., 2020; Ishida et al., 2021), and some of these studies have suggested that hyperexcitation of Emx1-Cre^+^ forebrain excitatory neurons contribute to premature mortality in epilepsy (SUDEP)(Boillot et al., 2014; Soh et al., 2014; Bunton-Stasyshyn et al., 2019; Klofas et al., 2020; Ishida et al., 2021).

Despite the assumption that the Emx1-promoter is specific to forebrain excitatory neurons, earlier studies reported extracerebral expression of Emx1. Endogenous Emx1 is detected in the urogenital tract, branchial pouches, apical ectodermal ridges, and some cervical ganglia of the developing mouse embryo (Briata et al., 1996), and this expression pattern is reflected in the Emx1-IRES-Cre mouse line (Gorski et al., 2002; Zhou et al., 2021). Thus when used in the developmental Cre-loxP approach, the functions of the targeted gene expressed in these peripheral cells can be also modulated and might affect the overall phenotype, if the gene of interest is expressed in these cells. However, the physiological roles of extracranial peripheral Emx1-cre^+^ cells are not well characterized in the postnatal mouse.

In this report, we identified that peripheral Emx1-Cre^+^ cells in the sensory vagus nerve directly affect cardiorespiratory regulation. Our anatomical study using a fluorescence reporter identified Emx1-Cre expression in their afferent vagal nerve fibers innervating the dorsal medulla as well as cell bodies in nodose ganglia and their potential projection sites at the carotid bifurcation. Optogenetic stimulation of the Emx1-Cre^+^ afferent fibers in acutely prepared brainstem slices *in vitro* confirmed functional synaptic input onto primary neurons within the nucleus tractus solitarius (NTS), and triggered parasympathetic bradycardia in anesthetized mouse preparations *in vivo*. The ascending projection from the NTS neuron has a complex influence on forebrain functions involved in somatic and autonomic systems. These results raise caution when interpreting behavioral and neurophysiological phenotypes of transgenic mice using the Emx1-IRES-Cre line.

## Material and Methods

### Animals

All animal experiments were conducted under the protocol approved by the institutional IACUC and conducted in accordance with the *Guide* of AAALAC. Emx1-IRES-Cre (Emx1-Cre, JAX stock number: 005628), floxed-ChR2 (Ai32, JAX stock number: 012569), and floxed-tdtomato (Ai9, JAX stock number: 007909) lines were obtained from Jackson laboratory. These animals were crossed and maintained in the vivarium with standard mouse chow. Most of the experiments were conducted on animals aged P25-90, except for the optogenetic studies in which young adult animals (~P40) were used because older animals lacked consistent ChR2 expression. We initially conducted studies using male and female mice separately. However, because we did not detect major differences in Cre expression patterns, both male and female mice data were pooled.

### Fluorescence imaging

Adult Emx1-Cre:tdtomato mice were transcardially perfused with PBS followed by 4% formalin. Brains were extracted and kept in 4% formalin overnight at 4°C, then transferred to 30% sucrose solution until the brain sinks. Fixed brains were frozen in OTC, cut in 20 μm sagittal or coronal sections, and mounted on glass slides. After washing in PBS, sections were coverslipped with a mounting medium (Prolong gold antifade, ThermoFishher) and analyzed on a Keyence BZ-X microscope. In the higher magnification image shown in **Figure 1F**, the image was deblurred using a deconvolution algorithm.

**Figure 1.**
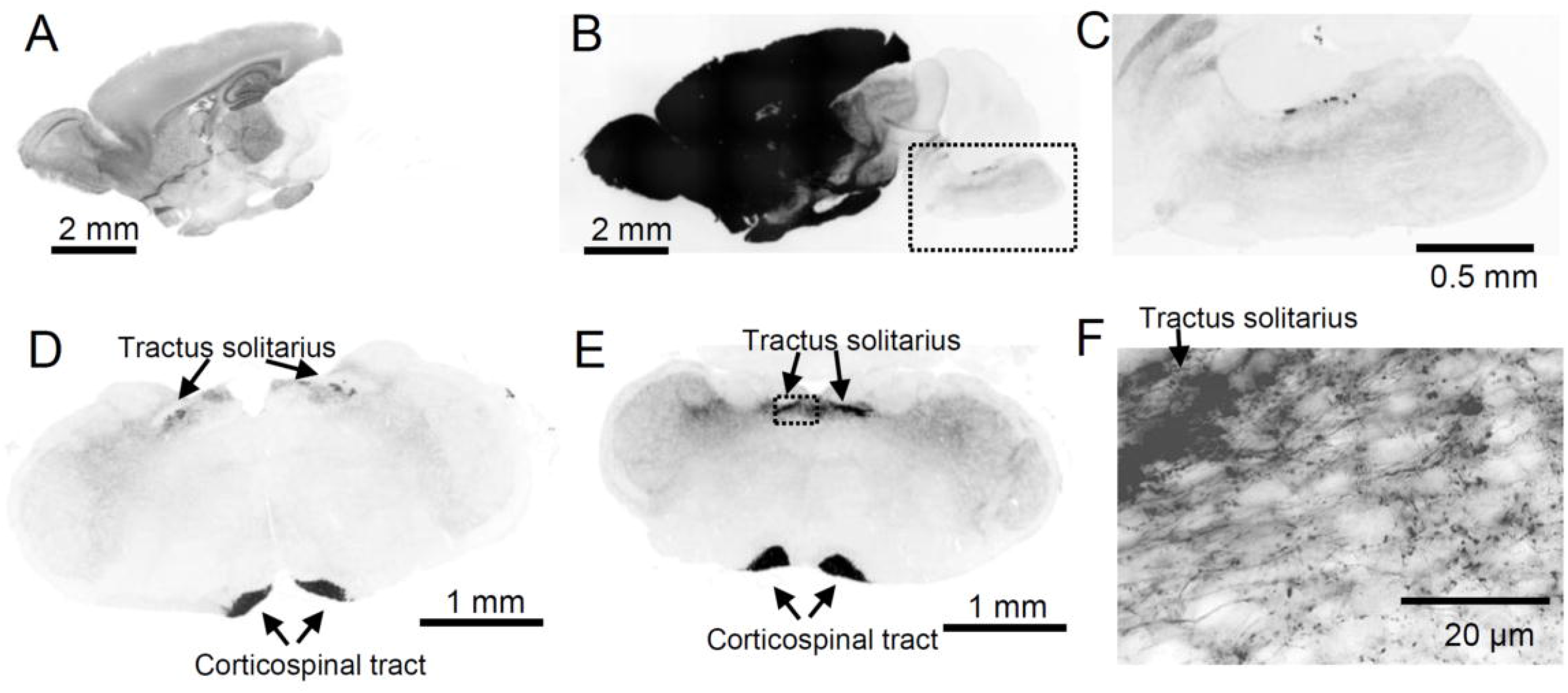
Characterization of Emx1-tdtomato signals in the brain. The fluorescence signal is inverted and shown in black color. **A-C** A midsagittal section shows strong fluorescence in forebrain structures. The overexposed image in B shows minor but significant signals present in hindbrain regions. The box in **B** is enlarged in **C. D-F** Coronal sections of Rostral (**D**) and caudal (**E**) medulla. Fluorescence signals were present in the tractus solitarius and corticospinal tract. The box in E is shown enlarged in F. Dense fluorescence is present in the tractus solitarius, and labeled single axonal fibers with varicosities are detected in the NTS field.

The nodose ganglion and the branchpoint of the carotid artery including superior cervical ganglion were also extracted from formalin-fixed mice and incubated in 4% formalin overnight at 4°C. A whole-mount tissue was placed on a glass slide, and differential interference contrast (DIC) and fluorescence images were acquired. In some experiments, the isolated ganglia were cut in 10 μm frozen sections to characterize their microstructures. In **Figure 2B**, the background fluorescence signal was eliminated by adjusting the threshold.

**Figure 2.**
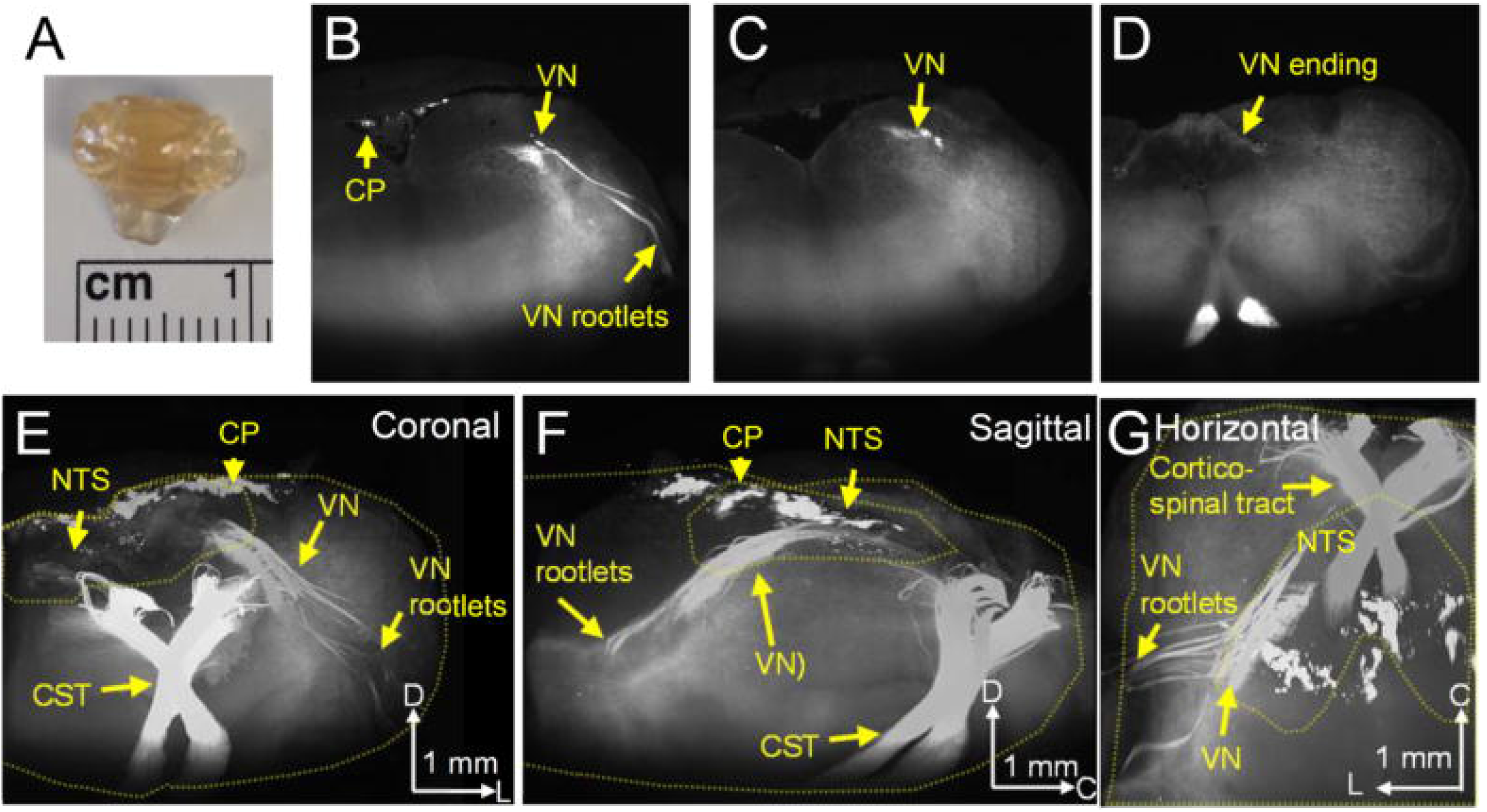
3D imaging of Emx1-tdtomato signals in the brainstem. The formalin-fixed brainstem was optically cleared and scanned on a light-sheet microscope. A. cleared brainstem preparation. B-D. Coronal serial images of Emx1-tdtomato signals in the left half of the brainstem. At the rostral level, the Emx1-tdtomato signal is detected in the vagus nerve rootlets of the brainstem (B), traveling the dorsal medulla at the caudal level (C), and terminates within the NTS at the caudal end (**D**). The full scanned image is available as **Supplementary video 1. E** G. 3D reconstructed of Emx1-tdtomato signals presented in coronal (**E**), sagittal (**F**), and horizontal view (G). The Emx1-tdtomato^+^ vagus nerve traverses the medulla. Outlines of brainstem tissue and NTS are depicted by yellow dashed lines. The full image is available as **Supplementary video 2**. Arrowheads in the scale bar indicate the axis, V: ventral, L: lateral, C: caudal. Abbreviation: VN: vagus nerve, CP Choroid plexus, NTS: Nucleus tractus solitarius, CST: Corticospinal tract

In some studies, extracted tissues were optically cleared using the EZ clear method (Hsu et al., 2022). In brief, PFA perfused tissues were incubated in 50% tetrahydrofuran overnight, washed in distilled water for 4 hours, and then further incubated in 80% Nycodenz/7 M Urea solution >24 hours to adjust tissue refractory index. The cleared tissues were imaged on a Nikon TE2000-S fluorescence microscope with a 4x objective or Zeiss Lightsheet Z1 microscope with a 5x objective and analyzed using the ImageJ program.

### *In situ* hybridization

The mRNAs in brain sections were detected using probe sets and buffers purchased from Molecular instruments. The probes used in this study are for ChAT, TH, and NPY each consisting of 20 DNA probe sets. The formalin-fixed brain sections (16-20 μm thickness) were mounted on glass slides and dehydrated by serially incubated in graded concentrations of ethanol (50%, 75%, 100%, 5 minutes each). The tissue was air-dried, treated with proteinase K for 10 minutes at 37°C, washed with 5xSSCT, preincubated in hybridization buffers for 30 minutes at 37°C, and then incubated with the DNA probes at 37°C overnight. The tissue sections were washed with the wash buffers, 5xSSCT, preincubated in the amplification buffer, and then incubated with the amplifiers overnight at room temperature. The tissues were washed with 5xSSCT, stained with DAPI (0.1mg/ml), and coverslipped with Prolong gold antifade (ThermoFishher). Fluorescence images were acquired as described above.

### Acute brainstem slice preparation

Acute brainstem slices were prepared as described previously (Aiba and Noebels, 2015). Briefly, mice were deeply anesthetized with a mixture of ketamine/xylazine (35 mg/75 mg per kg mouse, i.p.), transcardially perfused with dissection solution (110 mM N-Methyl-D-glucamine HCl (pH 7.4), 6 mM MgSO_4_, 25 mM NaHCO_3_, 1.25 mM Na_2_HPO_4_, 3mM KCl, 10 mM glucose, 0.4 mM sodium ascorbate, saturated with 95% O_2_/5% CO_2_ gas), decapitated, the brain extracted, and 200 μm coronal slices were cut using a vibratome (Leica VT1200S). Slices were recovered in dissection solution at 33°C for 10 minutes, then transferred and kept in artificial cerebrospinal fluid (ACSF: 130 mM NaCl, 3 mM KCl, 25 mM NaHCO_3_, 1 mM MgSO_4_, 1.25 mM Na_2_HPO_4_, 10 mM glucose, 0.4 mM sodium ascorbate, saturated with 95% O_2_/5% CO_2_ gas) at room temperature until transferred to the experimental chamber.

### *In vitro* electrophysiology of nucleus tractus solitarius (NTS) neurons

Whole-cell recordings were made from visually identified NTS neurons. The recording chamber was continuously superfused with ACSF at 2.5-3.0 mm/min and maintained within 33-34°C. The recording pipette contained Cs-gluconate based internal solutions (in mM: 130 gluconic acid, 10 HEPES, 1 MgCl_2_, 8 NaCl, 10 BAPTA, 5 TEA-Cl, 5 QX314, 2 Mg-ATP, 0.3 Na-GTP, pH adjusted to 7.2 with CsOH). Evoked EPSCs were recorded at −70mV and 0mV after correcting for a 10 mV of measured liquid junction potential. EPSCs were evoked by photostimulation of surrounding tissue using a 200 μm diameter silicon tip LED (Thorlabs), delivered for 10 ms, at 20 Hz, 10 times. Stimulation strength was adjusted to obtain supraminimum responses and was typically within 0.5-1.5 mW. Signals were amplified with a Multiclamp 700B amplifier, digitized, and analyzed using pClamp10 software (Molecular instruments). The individual EPSCs were detected based on a template search and events >4 pA were included. The variability in EPSC onset (jitter) was calculated based on the time to the maximum slope of optogenetically evoked EPSCs in each recorded cell.

### Cardiorespiratory response to optogenetic vagal nerve stimulation in the anesthetized mouse

Emx1-IRES-Cre:ChR2 mice (~P40) were anesthetized with urethane (1.2-1.5 mg/kg, i.p.), allowing spontaneous respiration. Body temperature was maintained at 36.5-37.0°C using a feedback heating blanket and rectal thermistor. Oxygen saturation level (SpO_2_) was periodically tested from a depilated left thigh using the MouseOx sensor (Starr life science) and was always above 97.5% throughout the experiments. A pair of needle electrodes were inserted into the thoracic wall for electrocardiography, and a piezo disc was placed on the abdomen to detect breathing. Physiological signals were amplified using a Bioamp (ADI), digitized, and analyzed using the Labchart system (ADI), and were monitored throughout the experiments. Heart and respiratory rates were first calculated using the built-in cyclic measurement, followed by manually correction of detection error.

For dorsal medulla stimulation, mice were set on a stereotaxic frame in the prone position. After achieving a state of stable anesthesia, hair covering the dorsal neck was removed, the skin was cleansed, and a midline incision was made to expose the occipital bone. The neck muscles were carefully separated, and a part of the occipital bone and ligament was removed to expose the surface of the dorsal medulla. An LED fiber (1.25 mm tip size, 200 μm core diameter) connected with a spacer was placed over the dorsal brainstem surface and photostimulation was delivered using an LED power controller (LEDD1B, Thorlabs) and Master 8 (AMPI). The parameter used for stimulation was 20 Hz, 10 ms, and 400 pulses. Typical stimulation power was within 2.0-2.5 mW. This stimulation parameter was chosen to minimize the risk of depolarization block of stimulated nerves, while maintaining the time window to detect short term plasticity properties. The stimulation site was adjusted to obtain optimal responses. Once the baseline response was determined, the mouse was injected with atropine (1 mg/kg, i.p.) to block parasympathetic activity.

For peripheral photostimulation, mice were immobilized in the supine position. The hair covering the neck was depilated, the skin was cleansed, and a midline incision was made between the sternum and mandible. The fat tissue and muscles were separated by blunt dissection, and the left common carotid and vagus nerve were identified within the carotid sheath. Further dissection was made superiorly until the carotid bifurcation was identified. At this level, the carotid artery and vagus nerve were carefully dissociated. Preparations with significant bleeding were discarded as blood occluded the photostimulation. For optogenetic vagus nerve stimulation, a piece of parafilm was inserted under the vagus nerve to selectively photostimulate the isolated nerve. For photostimulation of the carotid bifurcation, photostimulation was directly applied to the exposed carotid bifurcation at a distance ~500 μm. Care was taken to avoid stretching or compressing the airway, nerves, and blood vessels. The photostimulation protocol and LED were the same as those used in the dorsal medulla study. In some mice, stimulation was tested twice in the same animals while altering the stimulation sites. In the control study, we confirmed that the same photostimulation of the peripheral vagus nerve or carotid bifurcation does not trigger any response in the mouse without ChR2 (n=4).

### Statistics

Data are presented as mean ± standard error and statistical methods are described in the text. Statistics were computed by the R program or Graphpad prism software.

### Drugs

Nycodenz was obtained from Accurate Chemical. Capsaicin was from Tocris. Atropine and other chemicals were purchased from Sigma Aldrich.

## Results

### Emx1-Cre is expressed in peripherally afferent vagal nerve fibers that terminate within the dorsal medulla brainstem

We characterized the distribution of Emx1-Cre expression by crossing Emx1-IRES-Cre with floxed-tdtomato mice (hereafter Emx1-tdtomato). Emx1-Cre driven tdtomato expression in the cerebral cortex was obvious due to the gross pink coloration of the tissues. The cortico-spinal tract was detected as a faint pink line running along the midline of the ventral brainstem. In sagittal sections, a strong fluorescence signal was obvious in forebrain structures (**Figure 1A**) as well as in the diencephalon due to corticofugal projections. We observed significant Emx1-tdtomato expression within the brainstem with circumscribed regions of expression seen in the dorsal pons/medulla (**Figure 1 B-C**).

In order to identify the origin of brainstem Emx1-Cre^+^ signals, we next analyzed a series of coronal sections. A strong signal was detected in the medial ventral area corresponding to the cortico-spinal tract (**Figure 1D&E**). Fluorescence signals were also detected in the dorsal medulla within the nucleus of tractus solitarius (NTS). In high magnification images, these were identified as a dense signal along tractus solitarius outgoing fine fibers with axonal varicosities (**Figure 1F**). The fluorescent nerve fibers were enriched within the NTS, while sparse axon fibers innervated the lateral structures involving the trigeminal nucleus.

The localized expression of Emx1-tdtomato^+^ nerves in the dorsal medulla suggests Emx1-Cre expression in the afferent vagus nerve. In fact, 3D imaging of the optically cleared brainstem (**Figure 2A**) revealed that the Emx1-tdtomato^+^ originate peripherally, originally detected as vagus nerve rootlets, which merge and traverse the brainstem caudal/medial/dorsal direction until terminating in the NTS where the nerve forms fine axonal fibers (**Figure 2B-G**, see **supplementary video 1&2** for full image). This result confirmed that the afferent vagus nerve contributes the majority of the dense Emx1-tdtomato^+^ nerve fibers in the dorsal medulla. In the same preparation, dense cortico-spinal nerve tracts were detected at the ventral medulla. These nerve bundles run the ventral end of the medulla, traverse the dorsal direction, and crossed at the level of the caudal end of NTS. We also detected the Emx1-tdtomato signal in subset of choroid plexus over the dorsal brainstem surface. The same Emx1-tdtomato expression pattern was detected in three cleared brainstem preparations.

The majority of the afferent vagus nerve originates from bipolar neurons residing in the nodose ganglion. In fact, the whole resected nodose ganglion showed tdtomato fluorescence (**Figure 3A-D**). The fluorescence signal was barely visible in the vagus nerve at low magnification, likely due to the high light refraction of the ensheathing myelin. Within the nodose ganglion, Emx1-tdtomato signals were detected in the soma as well as in the nerve fibers of ganglion cells (**Figure 3E**). We examined whether the vagus premotor nerves also contribute to the Emx1-tdtomato^+^ signal of the vagus nerve. Vagus motor nerve fibers are mostly contributed by cholinergic premotor neurons within the dorsal motor vagus nucleus and nucleus ambiguus. None of the cholinergic premotor neurons in these nuclei express Emx1-tdtomato (n=196 cells, **Figure 3F**). Thus Emx1-Cre is expressed selectively in the sensory vagus, but not in the premotor nerves.

**Figure 3.**
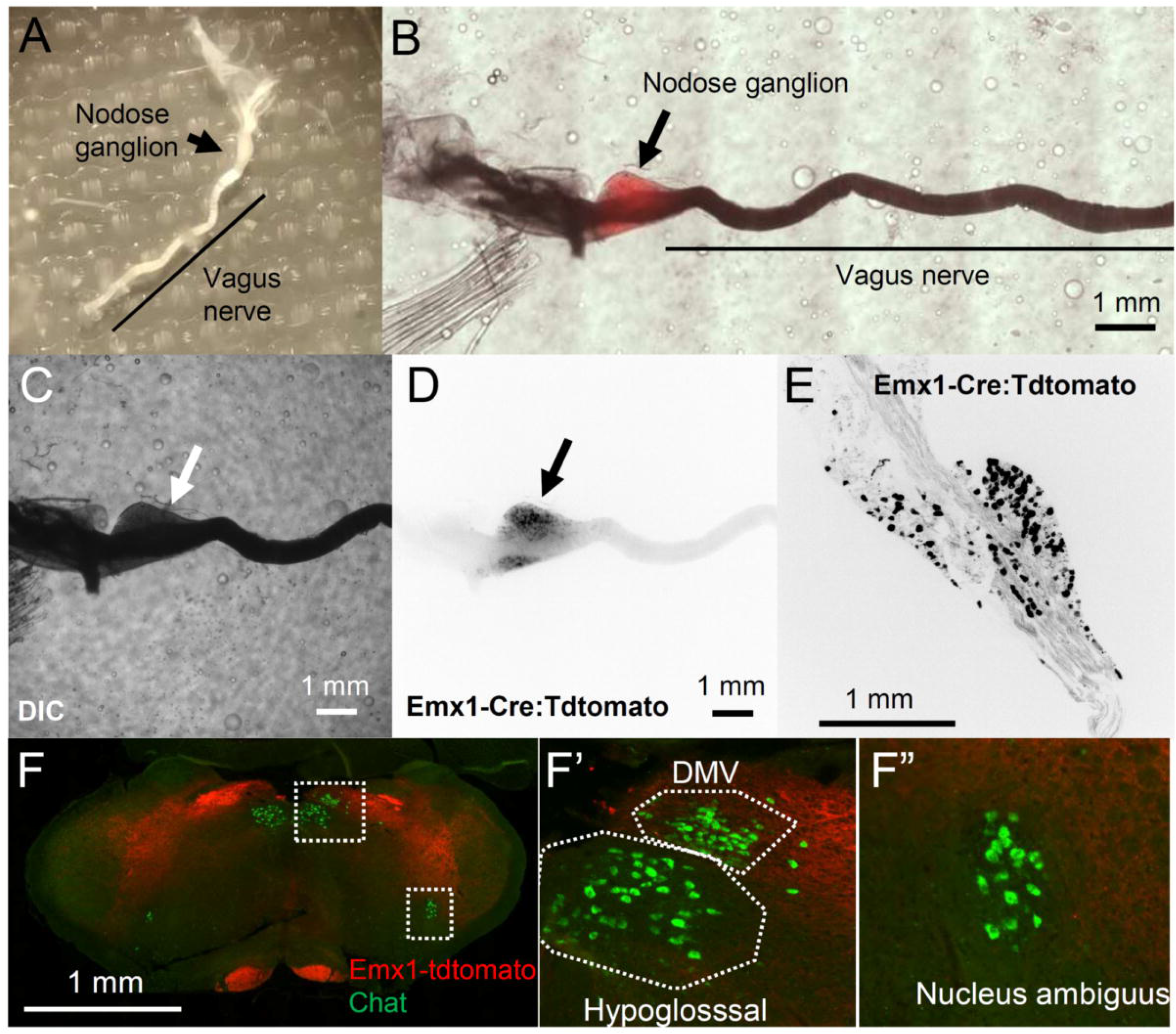
Characterization of Emx1-tdtomato signals in the nodose ganglion. A. Dissected nodose ganglion on the vagus nerve. B. Pseudocolor image merging, DIC (C), and tdtomato fluorescence (D) images of a whole-mount nodose ganglion. E. high magnification image of a sectioned nodose ganglion. Note fluorescence signals in D&E are inverted and shown in black. F. Emx1-tdtomato is not expressed in the Chat^+^ premotor neurons within the dorsal motor vagus nucleus (DMV) or Nucleus ambiguus neurons expressed Emx1-tdtomato.

In order to further characterize peripheral Emx1-Cre expression, we analyzed the Emx1-tdtomato signal in the neural tissue at the carotid bifurcation where the vagus nerve innervates the vascular sensory structures such as the carotid body, carotid sinus, as well as the superior cervical ganglion (SCG) (Wang et al., 2019). In the extracted whole-mount preparation (**Figure 4A**), blur Emx1-tdtomato fluorescence was identified at the carotid artery bifurcation and diffusely in the SCG (**Figure 4B&C**). Higher-resolution imaging of the optically cleared tissue detected Emx1-tdtomato^+^ nerve fibers at the carotid sinus of the internal carotid artery as well as in the SCG (**Figure 4D**). Emx1-tdtomato signals in the SCG were further analyzed in thin tissue sections. In this preparation, the Emx1-tdtomato signals are detected as fragments of nerve fibers and are not detected in the Tyrosine hydroxylase (TH) or Neuropeptide Y (NPY) positive sympathetic postganglionic cell bodies (**Figure 4E&F**). At the surface, some Emx1-tdtomato signals on the SCG surface colocalized with NPY signals. These results suggest the Emx1-tdtomato nerve fibers innervate the peripheral cardiovascular sensory sites.

**Figure 4.**
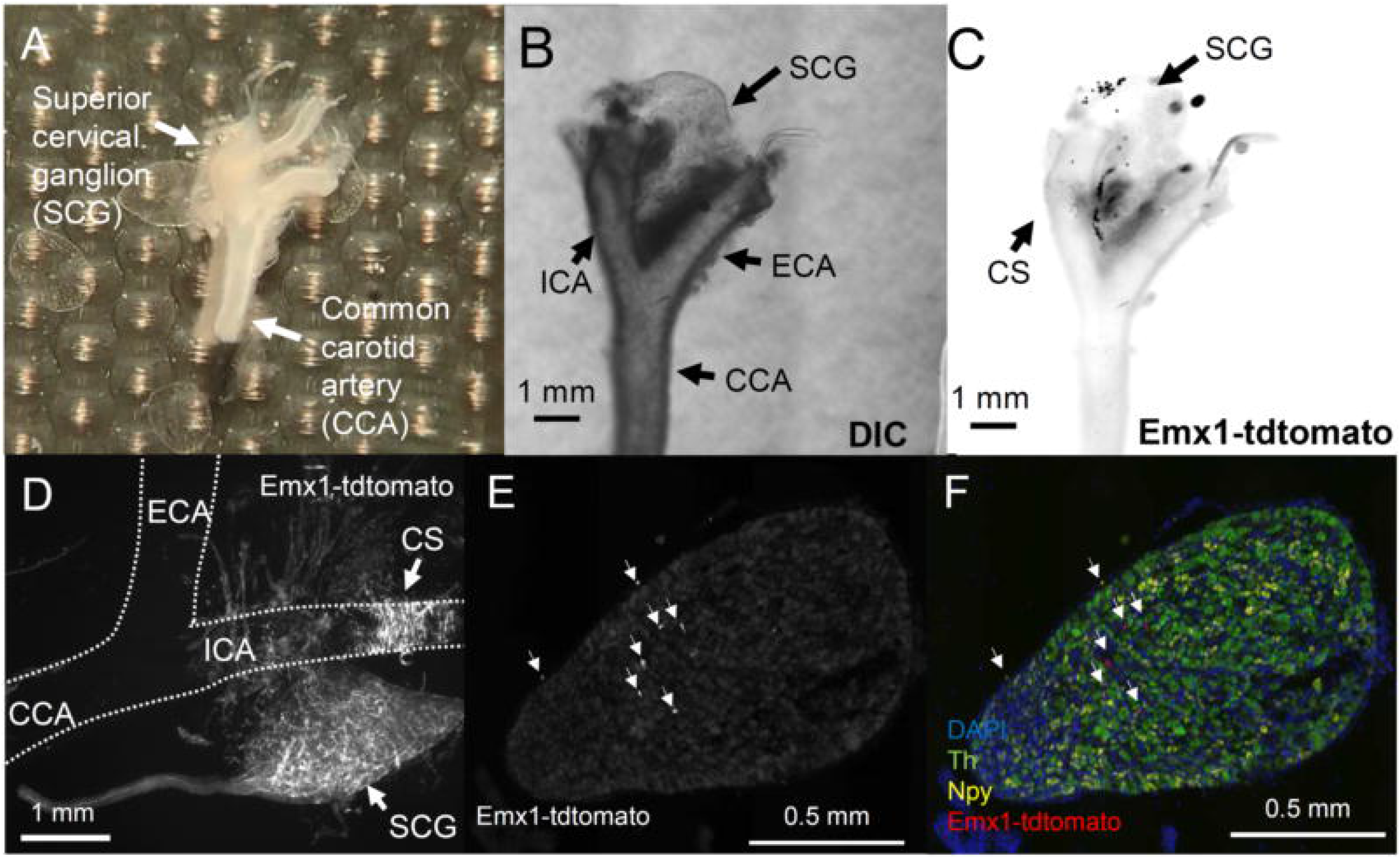
Characterization of Emx1-tdtomato signals at the carotid bifurcation. **A**. Dissected carotid artery and associated neural tissue, as well as the superior cervical ganglion (SCG). **B&C** A whole-mount image of dissected tissues in DIC (B) and Emx1-tdtomato fluorescence (C). Note fluorescence is shown in black in figure **C. D**. higher magnification image of Emx1-tdtomato fibers in a cleared carotid tissue and SCG. Fluorescence signals (in white) are detected at the carotid sinus (CS), SCG, and surrounding nerve fibers. The outline of the carotid artery is shown by the dashed line. E&F. Emx1-tdtomato signals in the SCG section. Emx1-tdtomato signals are only sparsely detected as pieces of nerve fibers in the SCG (arrowheads). Sympathetic preganglionic neurons are identified by *in situ* hybridization of tyrosine hydroxylase (Th) and Neuropeptide Y (Npy). E. Emx1-tdtomato only, F color image merged with Th (green) and Npy (yellow). CCA: common carotid, ICA: internal carotid artery, ECA: external carotid artery.

In addition to these neural tissues, the Emx1-tdtomato signal was also observed in the salivary glands and kidney (not shown); the latter is consistent with an earlier study that reported endogenous Emx1 expression in the urinary tract during development. Macroscopic observation did not detect robust expression of Emx1-tdtomato in isolated heart or lungs, however we do not exclude possible expression in their microstructures.

Together, our anatomical study revealed a significant expression of Cre-mediated recombination in extracerebral structures, namely the sensory vagus nerve of the postnatal Emx1-IRES-Cre mouse. Because of their neuronal gene expression and the relevance to autonomic dysfunctions and premature death reported in studies using Emx1-IRES-Cre mice, we further characterized the physiological functions of the Emx1-Cre^+^ vagus afferent nerve.

### Photostimulation of Emx1-Cre^+^ vagus nerve terminals excites neurons within the nucleus tractus solitarius (NTS) in vitro

In order to test the physiological functions of Emx1-Cre^+^ afferent vagus nerve, we crossed Emx1-IRES-Cre mice with floxed-channelrhodopsin (ChR2) and characterized synaptic function using acutely prepared brainstem slices containing the NTS *In vitro*. Whole-cell recordings were made from the visually identified NTS neurons which receive monosynaptic input from the afferent vagal nerves (tractus solitarius, **Figure 5A**). Trains of photo-stimulations (10 ms, 20 Hz, 10 pulses) reliably triggered excitatory postsynaptic currents (EPSCs) in the recorded NTS neurons with a small variability in latency (0.65 ± 0.51 ms, n=36, **Figure 5A inset**), a characteristic of the monosynaptic vagal afferent connection (Peters et al., 2011). The evoked EPSCs were detected in all recorded NTS neurons (n=36 cells, 4 mice), suggesting that the majority, if not all, afferent vagus nerve fibers that synapse onto NTS neurons express Emx1-Cre during development. No responses were detected when photostimulation was tested in mice without ChR2 (10 cells, 3 mice).

**Figure 5.**
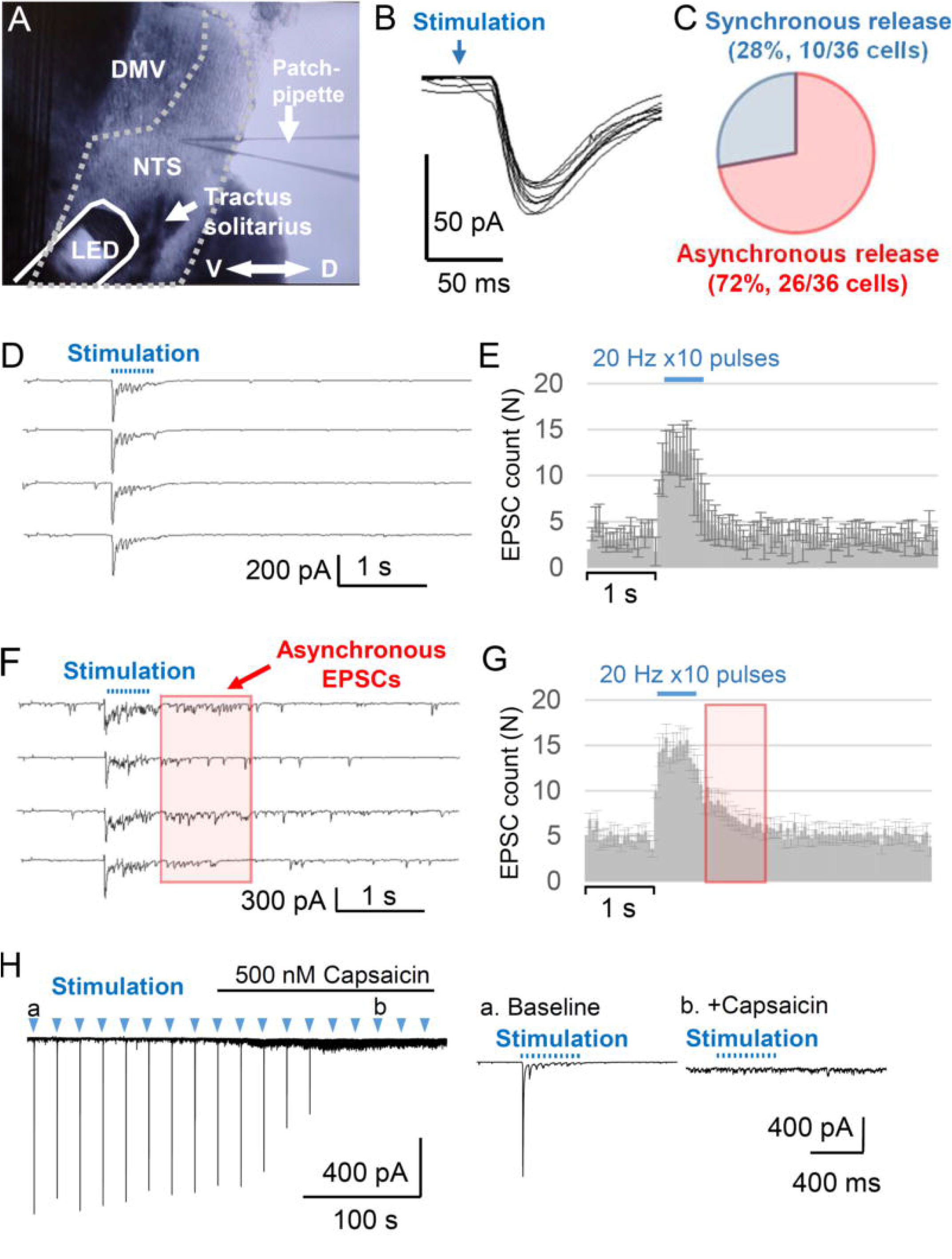
Optogenetic stimulation of Emx1-Cre^+^ nerve fibers *in vitro*. Vagus afferent nerve (tractus solitarius) was stimulated by a LED located over the brainstem slice (A). The border of NTS is indicated by the dashed line. Scale: 200 μm, axis D: dorsal, V: ventral. **C-G**. Photostimulation evoked either synchronous (D&F) or asynchronous EPSCs (F&G) in all recorded NTS neurons (n=36) with little variability in the onset time (B). C. 28% (10/36) of NTS neurons received synchronous, whereas 72% (26/36) of neurons received asynchronous EPSCs. **D&F** representative EPSC traces and **E&G** Histograms of mean EPSC (50 ms bins). Note the elevation of EPSC frequency following optogenetic afferent stimulation in E. values are mean ± standard error. H. Capsaicin effect on the NTS neurons receiving asynchronous EPSC. Capsaicin depolarizes the TRPV1^+^ vagus nerve terminals, resulting in increased sEPSC and block of optogenetically evoked EPSC. The inset shows responses to optogenetic stimulation during baseline (a) and after exposure to capsaicin (b).

The vagus afferent nerves are comprised of TRPV1^+^ and TRPV1^-^ fibers as characterized by the presence or absence of asynchronous presynaptic glutamate release, respectively (Peters et al., 2010, 2011), and each NTS neuron receives only one of either type of afferent input. We grouped asynchronous and synchronous release populations based on the presence of increased sEPSCs during the 1 s post-stimulation period (p<0.001, two-way ANOVA Mixed effect model). With this criterion, we identified cell groups with synchronous (28%, 10/36 cells **Figure 5B&C**) and asynchronous release (72%, 26/36 cells **Figure 5D&E**).

The TRPV^+^ vagus afferent nerve terminals are uniquely sensitive to capsaicin which depolarizes the afferent nerve terminals and blocks evoked synchronous glutamate release (Peters 2011). In our recording, bath application of capsaicin (500 nM) significantly enhanced the sEPSC, while at the same time, blocking evoked EPSCs in NTS cells receiving the TRPV1^+^ asynchronous release (**Figure 5F**). These results further indicate the Emx1-Cre^+^ presynaptic terminals showed properties consistent with the vagus afferent nerve. While we cannot exclude the contribution of Emx1-Cre^+^ cells in forebrain excitatory neurons (see Discussion), our results indicate that Emx1-Cre^+^ vagal afferent fiber populations functionally contribute to the brainstem autonomic circuits within the NTS.

### Optogenetic stimulation of Emx1-Cre^+^ structures acutely modulates cardiorespiratory function in the anesthetized mouse

Finally, we examined the physiological function of the Emx1-Cre^+^ afferent nerve *in vivo* using a urethane anesthetized mouse preparation (Emx1-ChR2^+/-^). Mice were anesthetized, an incision was made on the dorsal neck, and the left carotid artery and vagus nerve were isolated by blunt dissection. A LED was positioned to target the isolated vagus nerve at ~500 μm distance to avoid physical contact. Optogenetic stimulation (20 Hz, 20 s) of the isolated left vagus nerve evoked a fast transient bradycardia (33 ± 11% decrease from baseline, p<0.05, n=6, paired *t*-test) together with respiratory depression (67 ± 9% decrease from baseline, p<0.01, n=6, paired t-test **Figure 6B-D**). Optogenetic stimulation was also applied to the carotid bifurcation where Emx1-Cre^+^ nerve fibers were identified (**Figure 4**). Stimulation of the carotid bifurcation site triggered bradycardia (35 ± 13% decrease from baseline, p<0.05, n=5 trials, paired t-test) as well as respiratory depression (65 ± 11% decrease from baseline, p<0.05, n=5 trials, paired t-test, **Figure 6E-F**). Thus these peripheral Emx1-Cre^+^nerve fibers are involved in cardiorespiratory regulation.

**Figure 6.**
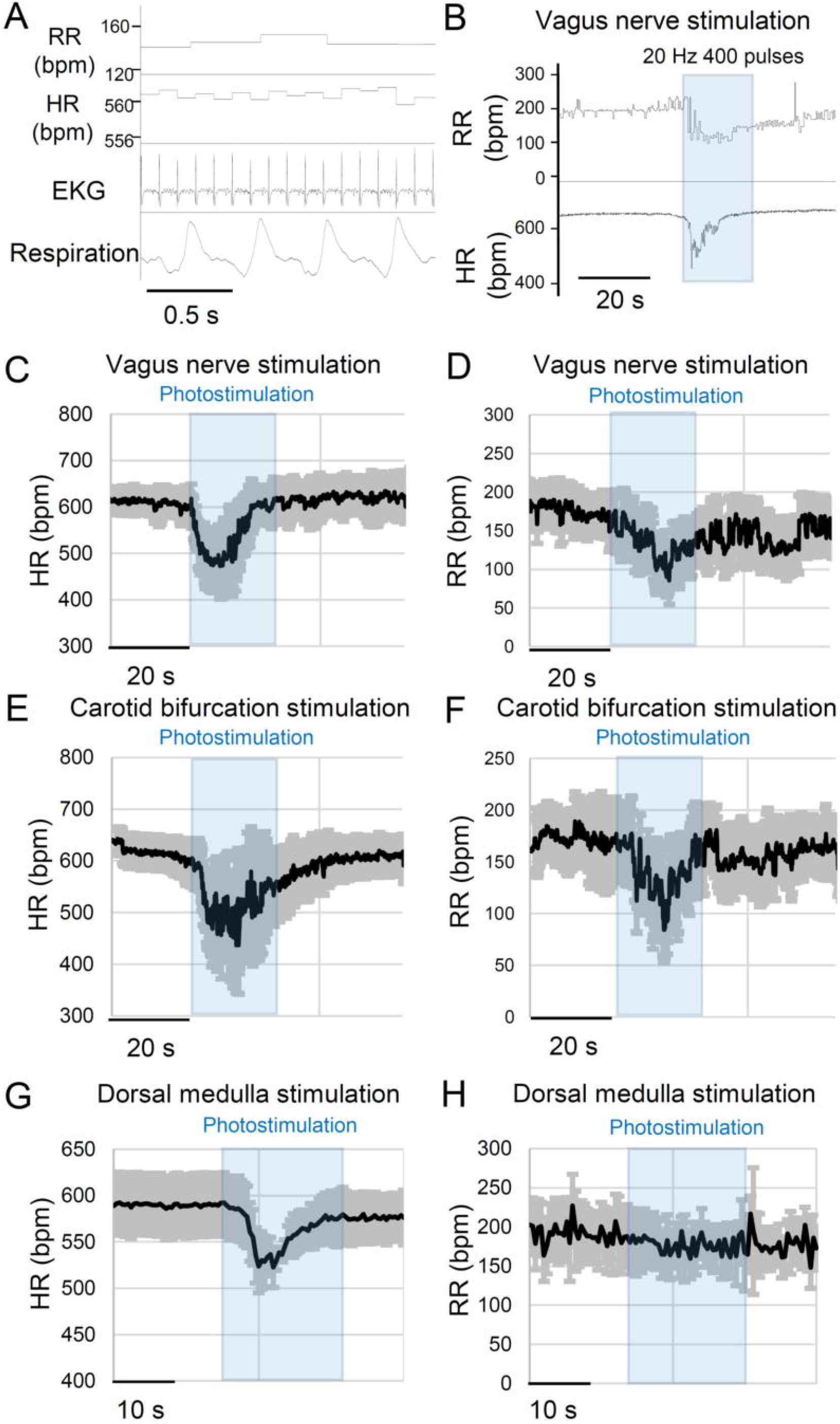
Optogenetic stimulation of Emx1-Cre^+^ nerve fibers *in vivo* urethane anesthetized mouse. A. EKG and respiratory signals were converted to heart rate (HR) and respiratory rate (RR). B. Example of respiratory and cardiac depression response to vagus nerve photostimulation. **C&D** Cardiorespiratory response to vagus nerve photostimulation. In these recordings, n=6 responses from 5 mice. E&F. Cardiorespiratory responses to photostimulation of the carotid bifurcation. n=5 responses from 5 mice. **G&H** Cardiorespiratory response to photostimulation of the dorsal medulla. An LED was horizontally positioned over the exposed dorsal medulla surface. A spacer was used to prevent direct contact. Stimulation consistently depressed heart rate while changes in respiration tone were variable. n=5 responses from 4 mice.

Finally, an additional optogenetic simulation study was conducted on the dorsal medulla of a urethane anesthetized mouse. The dorsal medulla surface was exposed by removing overlaying neck muscles and dura. Optogenetic stimulation (20 Hz, 20 s) of the dorsal medulla evoked fast bradycardia (14 ± 3% decrease from baseline, p<0.05, n=5, paired t-test) with an insignificant change in the respiration rate (**Figure 6G&H**). The weaker response seen in brainstem photostimulation may be due to co-activation of antagonizing pathways, a high light reflection property of the brainstem surface, and/or different position of the mouse under study (i.e. supine vs prone)

Together, these *in vivo* results indicate that peripheral Emx1-Cre positive cells are involved in central autonomic cardiac and respiratory regulation. Genetic manipulation of their excitability may alter the autonomic circuit stability.

## Discussion

In this study, we characterized the central afferent vagal nerve expressing Emx1-IRES-Cre, a Cre diver line commonly used to investigate the impact of forebrain excitatory neurons on neurological phenotypes. Consistent with previous studies, we found Emx1-Cre expression in peripheral organs, including the nodose ganglion whose parent cell bodies give rise to the vagus nerves detected in the dorsal medulla as afferent nerve terminals. Putative vagus nerve fibers were also detected at the carotid bifurcation. Our *in vitro* and *in vivo* electrophysiological characterization suggest that these peripheral Emx1-Cre^+^ structures contribute to cardiorespiratory regulation. The Emx1-Cre^+^ afferent vagus nerve likely play a key role, while Emx1-Cre^+^ may also be expressed in other peripheral cells and contribute to autonomic cardiorespiratory regulation. Thus, even when the phenotype of an Emx1-IRES-Cre based mutant mouse recapitulates a specific genetic disease, the phenotype cannot be solely attributed to forebrain excitatory neurons if the gene of interest is also expressed in the peripheral sites.

Endogenous Emx1 expression in the dorsal telencephalon has been detected as early as E9.5 (Simeone et al., 1992), thus the Emx1-IRES-Cre line is useful for characterization of neonate/pediatric genetic diseases which arise during prenatal circuit maldevelopment. However, Emx1 is also detected in peripheral tissues including branchial pouches, kidneys, developing limb appendages, and autonomic ganglia of developing mouse embryo (Gorski et al., 2002). We were unable to detect endogenous Emx1 transcripts in adult organs (data not shown), and thus these peripheral expressions may be only transient during development; however, Cre-loxP genomic recombination permanently alters the target gene expression in these structures. Our results confirmed that the embryonic Emx1 expression patterns are carried over into the postnatal adult mouse using Emx1-IRES-Cre:tdtomato reporter line.

Our anatomical and physiological studies suggest that Emx1-Cre is predominantly expressed in the parasympathetic sensory system whereas no Emx1-tdtomato^+^ cells were found within the SCG, which supplies the major sympathetic innervation to the head and neck (**Figure 3**). These differential expression patterns may reflect the role of Emx1 in the distinct developmental pathways; Most of the nodose ganglion neurons differentiate from the epibranchial placodes, while the SCG cells are derived from the neural crest (Baker and Schlosser, 2005; Huber, 2006; Streit, 2008). In addition to the vagus nerve, an earlier study suggests Emx1 expression in the other cranial nerves (Gorski et al., 2002). Thus, Emx1-Cre may be also expressed in other autonomic nerves, such as the glossopharyngeal nerve, and could contribute to the peripheral Emx1-tdtomato signals and cardiorespiratory responses observed in this study.

The present study focused on the contribution of peripherally innervated Emx1^+^ nerves. However, our study does not necessarily exclude a central autonomic contribution of Emx1-Cre^+^ forebrain excitatory neurons. Cortical structures such as the insular, anterior cingulate, and part of entorhinal cortices modulate autonomic functions (Cechetto, 2014). While the majority of cortical-brainstem pathways are polysynaptic, rare monosynaptic connections were identified between the motor/sensory cortex and NTS (Wang et al., 2015). Subcortical structures such as the amygdala have a strong influence on cardiorespiratory tone. While the main autonomic efferent neurons of the amygdala are the GABAergic neurons in the central amygdala which do not express Emx1-IRES-Cre, the excitatory neurons in the basolateral amygdala could contribute to brainstem signals. The bed nucleus of stria terminalis is another autonomic subcortical nucleus containing excitatory projection neurons, however Emx1-IRES-Cre is not expressed in these neurons (Gorski et al., 2002).

This study focused on the vagus nerve system because of its relevance to SUDEP studies. These cells share neuronal genes expressed in central excitatory neurons such as Kcna1, Kcnq2, Kcnq3, Scn1a, Scn8a (Glazebrook et al., 2002; Glasscock et al., 2012; Kollarik et al., 2018; Kupari et al., 2019; Sun et al., 2019), genes that are relevant to epilepsy. Comprehensive gene expression of vagal sensory neurons in the nodose ganglion has been identified by transcriptomic analysis (Kupari et al., 2019). On the other hand, omics studies could fail to reveal the enrichment of genes such as Kcnq2 and Kv1.6, despite their potential roles in shaping axonal excitability (Glazebrook et al., 2002; Sun et al., 2019). Other genes implicated in genetic neurological diseases may also be present at low abundance in vagal neurons, and modulation of their expression may affect central autonomic regulation.

Our study found that optogenetic stimulation of the vagus afferent nerve activates the majority of NTS neurons *in vitro* and stimulation of the vagus nerve *in vivo* induces a transient cardio-depression in urethane anesthetized mice. Both of these responses were transient in nature; optogenetically evoked EPSC trains showed a progressive decrease in their amplitude (**Figure 5**) which is consistent with the response evoked by electrical stimulation *in vitro* (Peters et al., 2010). Correlatively, peripheral optogenetic stimulation *in vivo* evoked bradycardia and bradypnea responses typically peaked within 10 seconds and then gradually recovered even when photostimulation was continuously delivered (**Figure 4**). Emx1-cre^+^ circuits may be normally involved in the rapidly sensitized reflex circuits, and sustained bradycardia seen in the pathological situation may require recruitment of additional neuromodulatory pathways such as tissue hypoxia (Kim et al., 2018) and metabotropic receptor mechanisms (Aiba and Noebels, 2019). While urethane anesthesia has a relatively small impact on the cardiorespiratory reflex system, it still blunts the dynamic range (Sapru and Krieger, 1979; Massey and Richerson, 2017). In this regard, the physiological properties of Emx1-Cre^+^ vagal nerves may be better appreciated in awake animal models.

This study focused on cardiorespiratory effects of the Emx1-Cre^+^ vagus nerve activation, however, afferent vagal nerves carry visceral interoceptive signals which are received by NTS neurons and are propagated not only to autonomic circuitries within the hypothalamus and brainstem, but also forebrain structures, both directly and indirectly (Naritoku et al., 1995; Kawai, 2018). These broad and bilateral central projections have been invoked to explain the therapeutic effects of vagus nerve stimulation across a range of neuropsychiatric disorders, including major depression and epilepsy. Thus modulation of afferent vagal nerve excitability by Emx1-Cre may also affect cortical hyperexcitability and neurobehavioral phenotypes.

In addition to Emx1, a similar experimental concern may be present in the parvalbumin (PV)-Cre mouse line, frequently used to target fast-spiking interneurons. In fact, PV is known to be expressed in the nodose and petrosal ganglia (Ichikawa and Helke, 1995; Sato et al., 2014), diaphragm (Jeckel-Neto et al., 1993), and renal tubules (Belge et al., 2007). The functional significance of these peripheral PV^+^ cells in autonomic physiology remains to be elucidated.

Together, the present study establishes the functional role of Emx1-Cre^+^ cells in the regulation of cardiorespiratory function. Caution is needed when interpreting alteration in systemic physiology resulting from developmental Cre-flox transgenic mouse studies using this Cre driver line.

## Acknowledgments

We thank Drs Krishnan and Maheshwari for their helpful comments on this manuscript.

## Figure legends

**Supplementary video 1**

Images of Emx1-tdtomato brainstem. Video shows rostral to caudal direction.

**Supplementary video 2**

3D reconstructed Emx1-tdtomato brainstem. Vagus nerve and emanating nerve fibers are seen at the dorsal level and dense corticospinal tracts are present at the ventral surface.

